# Linking Germline Telomere Removal to Global Programmed DNA Elimination in *Tetrahymena* Genome Differentiation

**DOI:** 10.1101/2025.09.21.677574

**Authors:** Kohei Nagao, Alix Lemoine, Tomoko Noto, Kazufumi Mochizuki

## Abstract

In the ciliate *Tetrahymena*, telomeres of the germline micronucleus (MIC) are removed and replaced by *de novo* telomere addition during somatic macronuclear (MAC) development. In this study, we investigated the kinetics and mechanism of the MIC telomere elimination. Comparison of the MIC and MAC genome sequences indicated that the MIC telomeres are excised from chromosomes as part of larger MIC-limited sequences (MLSs) through chromosomal breakage. We confirmed this using an optimized oligo-FISH protocol and found that their elimination occurs in parallel with other programmed DNA elimination processes. CRISPR-Cas9 disruption of a MLS-associated Chromosome Breakage Sequence (CBS) showed that elimination of the MLS was not blocked but instead led to loss of adjacent MAC-destined sequence (MDS), suggesting abnormal co-elimination. In biparental crosses of the CBS mutant, however, both MLS and MDS were retained, DNA elimination was broadly disrupted, and no viable progeny were produced. These findings indicate that chromosome breakage at MLS-associated CBSs is essential for the proper separation of MLSs and MDSs, ensuring correct DNA elimination and successful sexual progeny development. We propose that the MIC telomere elimination is subsumed within the broader process of programmed DNA elimination.

## Introduction

Telomeres are specialized structures found at the ends of linear chromosomes in eukaryotic cells. Highly proliferative cells, including germ cells, exhibit elevated telomerase activity to counteract telomere erosion, thereby preventing genome instability and cell cycle arrest (Blackburn et al., 2015; de Lange, 2018). In addition to telomere length regulation by telomerase, telomere dynamics are also influenced by cell type- and stage-specific chromatin environments (Arora et al., 2012; Cubiles et al., 2018; Schoeftner and Blasco, 2009; Zickler and Kleckner, 2023). However, how telomere regulatory mechanisms switch between different cellular environments remains poorly understood.

The ciliated protozoan *Tetrahymena* undergoes a drastic transition from germline to somatic telomeres: removal of germline telomeres and *de novo* formation of somatic telomeres. *Tetrahymena* harbors a diploid germline micronucleus (MIC) and a polyploid somatic macronucleus (MAC) within a single cytoplasm. The MAC is the site of transcriptional activity and is responsible for cellular gene expression, whereas the MIC remains transcriptionally inert during vegetative growth, functioning exclusively as the repository and transmitter of genetic information to subsequent sexual generations. Sexual reproduction is initiated when two cells of complementary mating types engage in conjugation (Figure 1A). During this process, the MIC undergoes meiosis and contributes to the formation of both a new MIC and a new MAC, while the parental MAC is selectively degraded and eliminated (Cole and Sugai, 2012).

**Figure 1.**
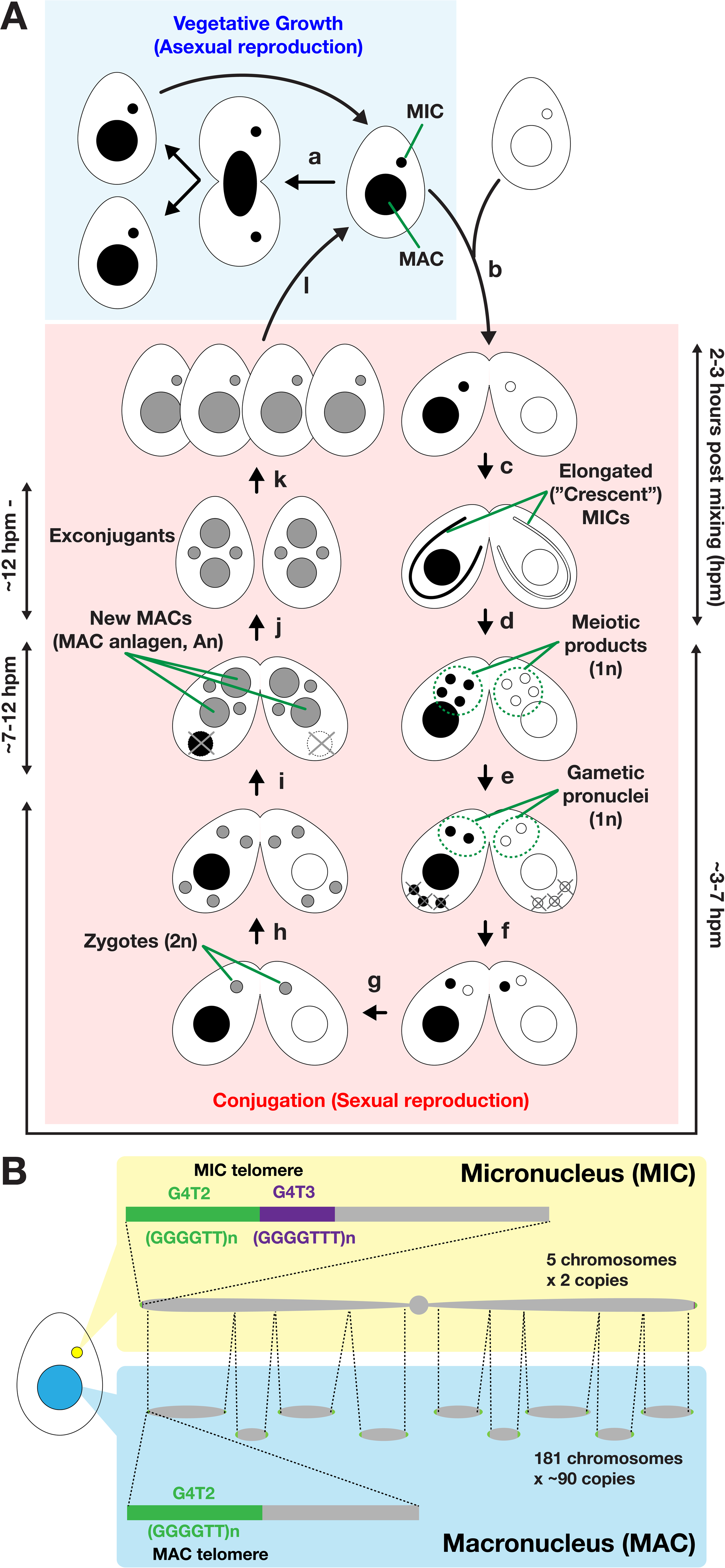
Life cycle and nuclear dimorphism of *Tetrahymena*. (A) Under nutrient-rich conditions, vegetative *Tetrahymena* cells proliferate by binary fission, with the macronucleus (MAC) and micronucleus (MIC) dividing independently (a). After prolonged starvation, two cells of complementary mating types fuse to initiate conjugation, which is the sexual reproduction process of *Tetrahymena* (b). During meiotic prophase, the MICs elongate into a crescent shape, in which chromosome arms are aligned in parallel, with centromeres and telomeres anchored at opposite ends (c). The MICs then undergo meiosis (d), and one meiotic product survives and divides mitotically to produce two gametic pronuclei (e). One of the two pronuclei of each side traverses the conjugation bridge (f) and fuses with the stationary pronucleus to form a diploid zygotic nucleus (g). The zygotic nucleus undergoes two mitotic divisions (h), producing four nuclei: two differentiate into MAC anlagen (An), while the other two remain as MICs; the parental MACs are degraded (i). The conjugating pair then separates to form exconjugants (j), which resume vegetative growth when nutrients are restored (k, l). (B) The MIC is diploid, with five chromosomes per haploid genome. MIC telomeres consist of distal G_4_T_2_ (5′-GGGGTT-3′) repeats (green) and proximal G_4_T_3_ (5′-GGGGTTT-3′) repeats (purple). In contrast, the MAC is polyploid (∼90 copies) and contains 181 chromosomes per haploid genome. MAC telomeres are composed exclusively of G_4_T_2_ repeats. The MAC is derived from the MIC during sexual reproduction, when chromosome breakage followed by *de novo* addition of G4T2 repeats generates MAC chromosomes. During this genome rearrangement process, MIC telomeres are thought to be eliminated from the developing MAC. Programmed DNA elimination of internal eliminated sequences (IESs) is not depicted.

The developmental remodeling of the MAC involves extensive genome reorganization (Figure 1B). Chromosome breakage at each conserved 15-bp Chromosome Breakage Sequence (CBS) fragments the five MIC chromosomes into 214 MAC chromosomes, of which 181 are retained (Hamilton et al., 2016; Yao et al., 1987). In parallel, programmed DNA elimination removes more than 12,000 internal eliminated sequences (IESs) via a small RNA–mediated heterochromatin assembly pathway, after which the flanking MAC-destined sequences (MDSs) are ligated (Chalker and Yao, 2011; Noto and Mochizuki, 2017). The restructured MAC chromosomes are amplified by endoreplication to approximately 90 copies each (Zhou et al., 2022).

While MAC telomeres consist of tandem G_4_T_2_ (5′-GGGGTT-3′) repeats, MIC telomeres contain an internal stretch of G_4_T_3_ (5′-GGGGTTT-3′) repeats followed by distal G4T2 repeats, as observed at all six MIC chromosomal ends studied to date (Kirk and Blackburn 1995). At all of these ends, the MIC-specific G_4_T_3_ telomeric repeat lies immediately adjacent to MIC-limited sequences (MLSs) (Kirk and Blackburn, 1995). This organization suggests that MIC telomeres are eliminated together with the adjacent MLSs, after which *de novo* addition of MAC-type telomeres occurs at the termini of the rearranged MAC chromosomes. However, the mechanism underlying MIC telomere elimination, as well as its relationship to other genome rearrangement processes, remains unresolved. In this study, we investigated the kinetics of MIC telomere and MLS elimination during MAC development, and the effect of genetic perturbation of chromosomal breakage on this process.

## Results

### MIC telomere elimination occurs in parallel to IES elimination in the new MAC

To study how MIC telomeres are eliminated during MAC development, we visualized the MIC-specific G_4_T_3_ telomere repeat using oligo-FISH and tracked its behavior during conjugation. We established an optimized protocol that included fixation, permeabilization, partial protein digestion with proteinase K, and removal of RNA-derived background with RNase A (see Materials and Methods).

To detect the G_4_T_3_ telomere repeat, we used an oligo DNA probe containing five tandem repeats of 5′-AAACCCC-3′, flanked by a sequence complementary to a fluorescently labeled secondary probe (Figure 2, top). Applied to wild-type *Tetrahymena* cells, the probe detected several foci in the nuclear periphery of the MICs in vegetative cells (Figure 2, Veg). At the early conjugation stage when MIC chromosomes align linearly, with centromeres and telomeres clustered at opposite poles (Loidl, 2021), the probe detected a few foci at the end of elongated MIC at meiotic prophase (Figure 2, 3 hpm). On the other hand, we only detected faint background level stainings in the MAC in these stages. These results confirm that our probe, together with the optimized preparation and hybridization conditions, specifically detects the MIC-specific G_4_T_3_ telomere repeat.

**Figure 2.**
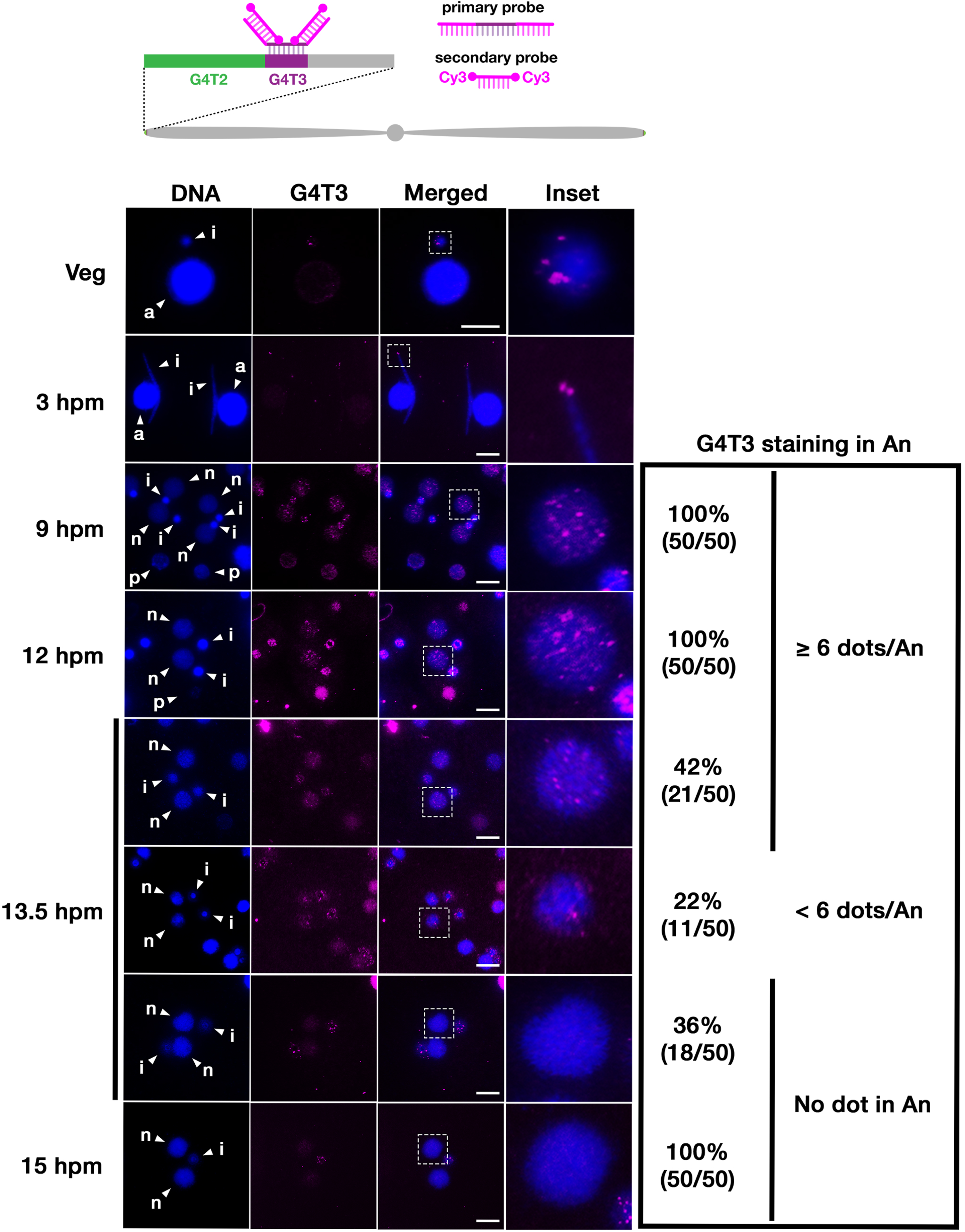
Elimination of MIC telomeres during MAC development. Vegetative (Veg) and conjugating wild-type cells at the indicated time points (hours post-mixing, hpm) were analyzed by oligo-FISH using an oligonucleotide probe complementary to the MIC-specific G_4_T_3_ repeat (magenta). DNA was counterstained with DAPI (blue). Insets show enlarged images of the regions indicated by dotted squares. The presence of the G_4_T_3_ FISH signal was quantified in 50 cells per time point and categorized according to the number of G_4_T_3_ dots per new MAC (An) (≥6 dots, <6 dots, or none). Arrowheads with “a”, “i”, “p”, and “n” indicate the MAC, the MIC, the parental MAC and the new MAC, respectively. Scale bars: 10 µm.

**Figure 3.**
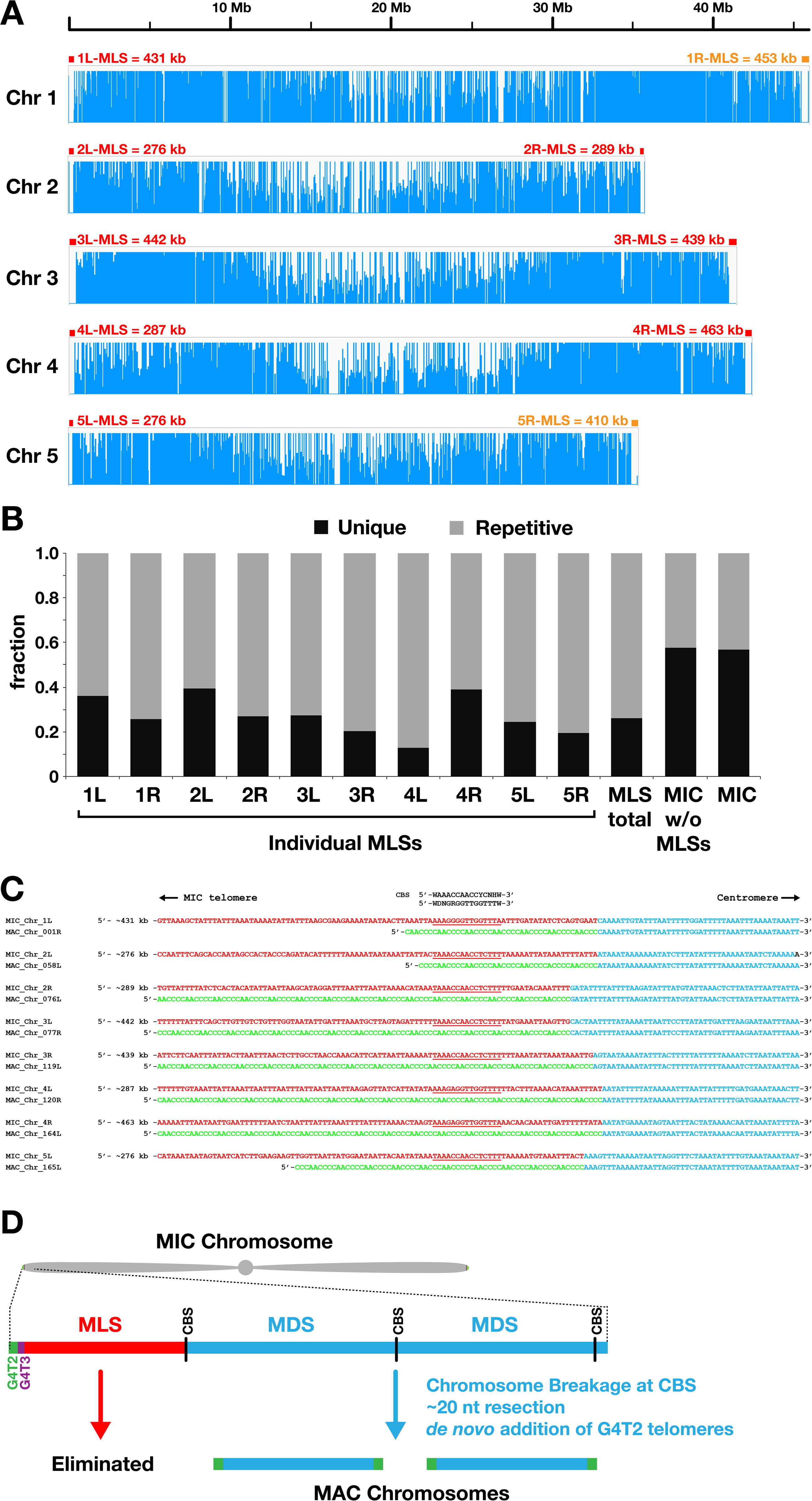
Identification of MIC-limited sequences at the ends of MIC chromosomes. (A) The MIC genome assembly (version 2021) and the MAC genome assembly (version 2025) were compared using Minimap2, and the coverage of each MIC chromosome sequence by the MAC genome sequence is shown as a histogram (blue). All MIC chromosome ends consist of 276–463 kb MIC-limited sequences (MLSs, marked by red or orange boxes) that are not represented in the MAC genome sequence. (B) The repetitiveness of each MLS was examined by mapping all possible 20-nt sequences from the MIC genome back to the same MIC genome and determining the fractions of each region that were covered (Unique) or not covered (Repetitive). (C) In eight of the ten MIC chromosome ends (red boxes in A), a chromosome breakage sequence (CBS) was found ∼20 nt distal to the boundary between MLSs (red) and MAC-destined sequences (MDSs, blue). The ends of the MDSs are directly linked to MAC telomeres (green). At the remaining MIC chromosome ends (orange boxes in A), no CBS was detected near the MLS–MDS border. (D) MIC telomeres are predicted to be separated from the MDSs by chromosome breakage at the CBS together with their associated MLSs and eliminated from the developing MAC. In parallel, the exposed MDS ends are subjected to ∼20 nt resection followed by de novo addition of G_4_T_2_ telomeres.

We then analyzed the dynamics of this repeat during new MAC development. Under our culture conditions, new MAC formation begins at ∼7 hpm, and most internal eliminated sequences (IESs) are removed between 10 and 18 hpm (Mutazono et al., 2019). G_4_T_3_ telomere foci were readily detected in new MACs at 9 and 12 hpm (Figure 2, arrowheads with “n”). For unknown reasons, we also detected a high level of staining in the parental MAC at these stages (Figure 2, arrowheads with “p”) which we believe represents nonspecific background, as it was also observed in the secondary probe-only control (data not shown). By 13.5 hpm, many new MACs showed either few (22%) or no (36%) foci, and by 15 hpm, no foci remained (Figure 2). This disappearance closely matched the elimination kinetics of the *Tlr1* element (Supplementary Figure S1), a repetitive transposable element embedded in IESs, and some other IESs (Mutazono et al., 2019). Together, these findings show that MIC telomere elimination occurs in parallel with IES elimination during new MAC development.

### MIC telomeres are likely co-eliminated with adjacent MIC-limited sequences through chromosomal breakage

Previous work showed that the MIC-specific G_4_T_3_ telomere repeat lies directly distal to certain MIC-limited sequences (MLSs) at least at six MIC chromosomal ends studied so far (Kirk and Blackburn, 1995). However, the exact configurations of MIC chromosome ends remain unclear.

To investigate this, we compared the latest assemblies of the MIC and MAC genomes (see Materials and Methods). Consistent with earlier findings, we confirmed that all MIC chromosome ends contain MLSs that are absent from the MAC genome. These MLSs range in size from 276 kb to 463 kb and lie immediately proximal to the G_4_T_3_ repeat (Figure 3A). We found that a large fraction of each MLS consists of repetitive sequences, ranging from 60.8% to 87.3% with an average of 73.8% (Figure 3B). This proportion is substantially higher than that observed in the total MIC genome, which is 43.0%. These suggest that MIC telomeres are eliminated together with their adjacent repeat-rich MLSs during MAC development.

We further observed that, at eight of the ten MIC chromosome ends, the conserved 15-nt Chromosome Breakage Sequence (CBS, 5′-WAAACCAACCYCNHW-3′) (Hamilton et al., 2006), which is essential and sufficient for inducing a chromosome break (Yao et al., 1990), is located ∼20 nt distal to the boundary between MLSs and adjacent MAC-destined sequences (MDSs) (Figure 3C). This ∼20 nt gap is consistent with previous reports of DNA end resection at chromosome breakage sites prior to *de novo* telomere addition during the formation of MAC chromosomes (Fan and Yao, 1996). Together, these findings suggest that, at least at eight MIC chromosome ends, MIC telomeres and their neighboring MLSs are separated from the rest of the chromosome by chromosome breakage and co-eliminated during MAC development (Figure 3D).

### MIC-limited sequences and MIC telomeres are eliminated with similar kinetics

To determine when MIC-limited sequences (MLSs) are eliminated during MAC development, we set out to visualize individual MLSs by oligo-FISH. Because MLSs are largely composed of repetitive sequences also found elsewhere in the MIC genome, we designed pools of oligo DNA probes uniquely complementary to a specific MLS (Supplementary Table S1; see also Materials and Methods).

We first targeted the MLS on the right arm of MIC chromosome 4 (4R-MLS), the longest MLS and one of the eight chromosomal ends containing a proximal Chromosome Breakage Sequence (CBS). A pool of 384 oligonucleotides (32 nt each), each with a 5’ extension complementary to a Cy3-labeled secondary probe, was synthesized and used for oligo-FISH. Similarly, another pool of 384 oligonucleotides specifically complementary to the adjacent MAC-destined sequence (4R-MDS), each with a 5’ extension complementary to a Cy5-labeled secondary probe, was synthesized and used for oligo-FISH (Figure 4, top).

**Figure 4.**
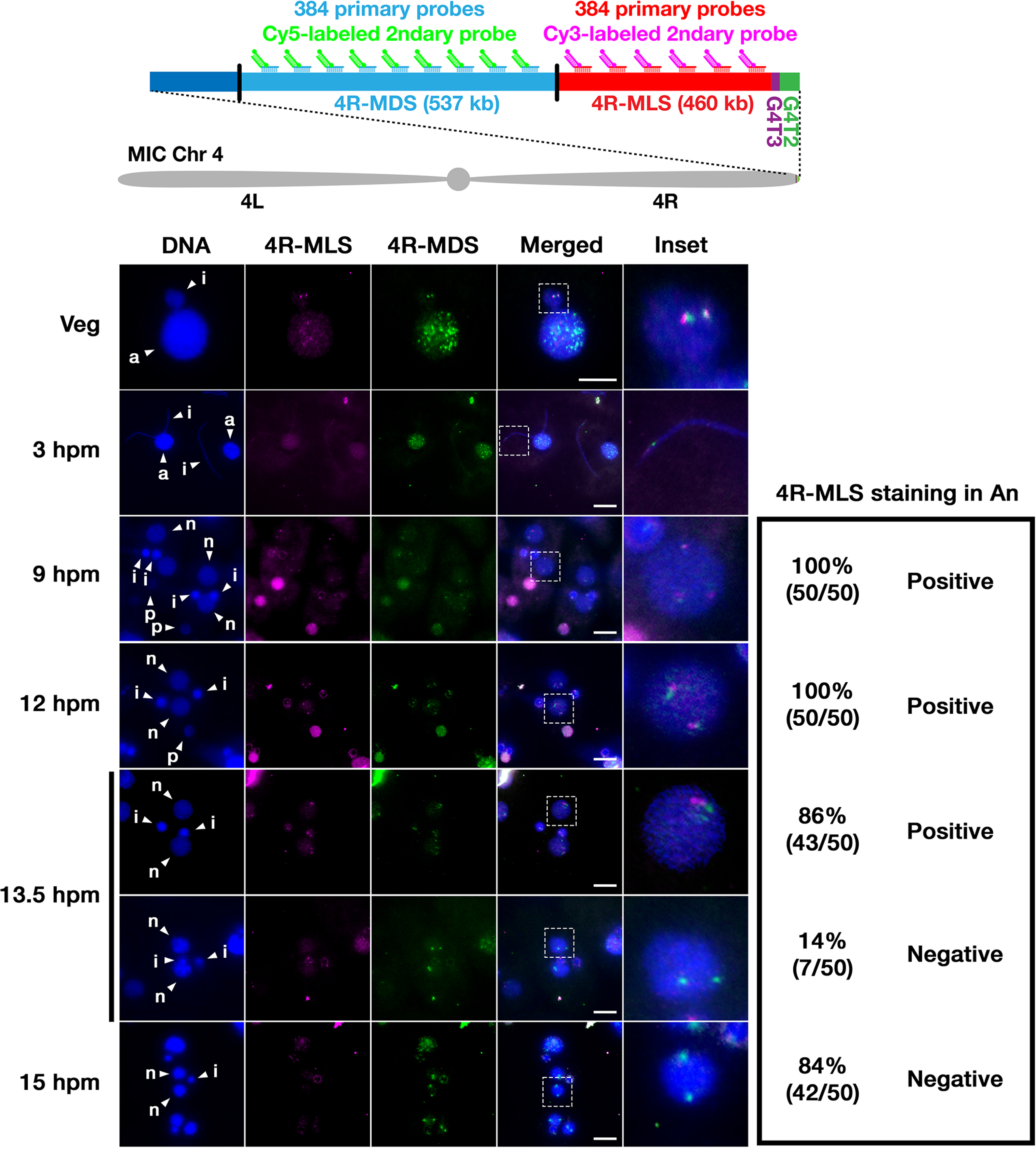
Elimination of 4R-MLS during MAC development in wild-type cells. Vegetative (Veg) and conjugating wild-type cells at the indicated time points (hours post-mixing, hpm) were analyzed by oligo-FISH using pool of oligonucleotide probes complementary to 4R-MLS (magenta) or 4R-MDS (green). DNA was counterstained with DAPI (blue). Insets show enlarged images of the regions indicated by dotted squares. The presence (Positive) or absence (Negative) of the 4R-MLS FISH signal in new MAC (An) in 50 cells per time point was examined. Arrowheads with “a”, “i”, “p”, and “n” indicate the MAC, the MIC, the parental MAC and the new MAC, respectively. Scale bars: 10 µm.

The 4R-MLS probes produced two distinct foci in vegetative MICs (Figure 4, Veg, magenta), which closely colocalized with foci marked by the 4R-MDS probes (Figure 4, Veg, green). At the early conjugation stage, both probes detected closely adjacent regions at the end of elongated MICs (Figure 4, 3 hpm). These results confirm that the probe pools specifically localize the 4R-MLS and 4R-MDS regions in the MIC.

We next analyzed the behavior of 4R-MLS and 4R-MDS during new MAC development. The 4R-MDS probe signal persisted in the new MAC from 9 hpm to 15 hpm (Figure 4, green). In contrast, the 4R-MLS signal began to disappear at 13.5 hpm and was largely undetectable by 15 hpm (Figure 4, magenta). A similar elimination pattern was observed for the MLS at the left arm of MIC chromosome 3 (3L-MLS, Supplementary Figure S2).

These results demonstrate that chromosome-end MLSs and the MIC-specific G_4_T_3_ telomere repeat are eliminated at similar times during new MAC development. This supports the idea that MIC telomeres and MLSs are co-eliminated following their separation from MIC chromosomes by chromosomal breakage.

### 4R-CBS mutation does not block elimination of 4R-MLS

To test the role of chromosomal breakage in eliminating MIC telomeres and MLSs, we disrupted the most distal Chromosome Breakage Sequence (CBS) at the right arm of MIC chromosome 4 (4R-CBS) using CRISPR–Cas9 mutagenesis (Figure 5A). A guide RNA (gRNA) complementary to the 4R-CBS locus was designed (Figure 5B, top). We established two homozygous 4R-CBS mutant lines from the wild-type B2086 strain and two from the wild-type CU428 strain (Figure 5B). All mutants lacked at least one nucleotide previously shown to be essential for inducing chromosome breakage in cis (Fan and Yao, 2000).

**Figure 5.**
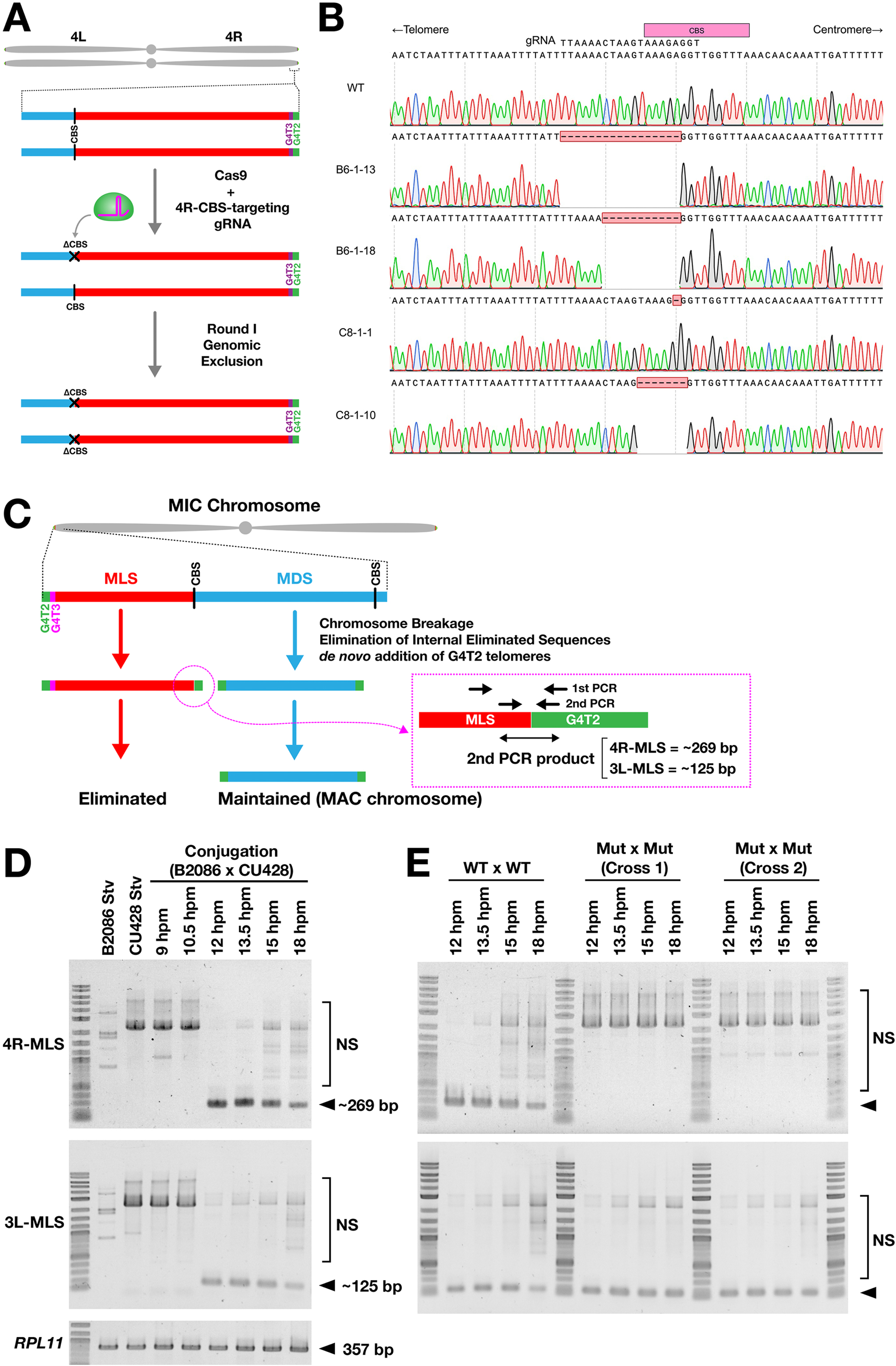
Production of mutants with deletions at 4R-CBS. (A) The most distal CBS at the right arm of MIC chromosome 4 (4R-CBS), which separates MLS (red) and MDS (blue), was targeted for mutagenesis using Cas9 and a guide RNA (gRNA) complementary to the 4R-CBS locus (see B, top). Heterozygous 4R-CBS mutants were obtained and then subjected to Round I genomic exclusion to generate homozygous 4R-CBS mutants. See Materials and Methods for detailed genetic procedures. (B) The 4R-CBS locus of two mutants derived from the B2086 strain (B6-1-13 and B6-1-18) and two mutants derived from the CU428 strain (C8-1-1 and C8-1-10) was compared with that of the wild-type strain (WT) by genomic PCR followed by Sanger sequencing. All 4R-CBS mutants carried deletions (red boxes) that removed essential conserved sequences of the CBS (pink box, top). (C) The MAC-type G4T2 telomere (green) is expected to be added *de novo* following chromosome breakage not only at the ends of MDSs (blue) but also at the proximal end of the MLS (red) prior to its elimination. A telomere-anchored PCR assay using nested PCR was designed to detect *de novo* telomere formation at MLS ends. (D) Genomic DNA was extracted from wild-type cells under starvation (Stv) and during conjugation at the indicated time points after mating induction (hours post-induction of mating, hpm) and used for the telomere-anchored PCR assay. The coding region of *RPL11* was also amplified from the same genomic DNA by single-step PCR as a control. Telomere-capped 4R-MLS and 3L-MLS ends are expected to produce PCR products of approximately 269 bp and 125 bp, respectively (marked with arrowheads). NS = non-specific amplification products. (E) Two independent crosses of 4R-CBS mutants were analyzed as (D).

We next sought to confirm that mutations at 4R-CBS indeed block chromosome breakage. Previous work showed that *de novo* telomere formation following chromosome breakage can be detected by telomere-anchored PCR not only at MDS ends but also at internal chromosomal fragments (eliminated minichromosomes, EMCs), which are not maintained and are lost during vegetative growth (Lin et al., 2016). Based on this approach, we designed a similar telomere-anchored PCR assay to detect *de novo* telomere formation at the ends of MLSs after chromosome breakage (Figure 5C).

In wild-type strains, telomere-capped MLS ends were undetectable during vegetative growth and throughout conjugation until 10.5 hpm, first became detectable at 12 hpm for both 4R-MLS and 3L-MLS, and then gradually declined by 18 hpm likely due to elimination of MLSs (Figure 5D). These results indicate that *de novo* telomere formation also occurs at MLS ends following chromosome breakage. We then examined the 4R-CBS mutants and found that de novo telomere formation was specifically blocked at the 4R-MLS end, while it occurred normally at 3L-MLS (Figure 5E). Together, these results confirm that mutations at 4R-CBS specifically block chromosome breakage at 4R-CBS.

We next examined how the 4R-CBS mutations affected elimination of 4R-MLS. Two of the homozygous 4R-CBS mutants were crossed with wild-type strains (WT × Mut crosses). In these crosses, new MACs inherit one wild-type and one mutant 4R-CBS locus. By 30 hpm, the 4R-MLS signal was almost entirely undetectable (Figure 6A) and started being eliminated by 13.5 hpm (Figure 6B). Therefore, 4R-MLS in WT × Mut crosses are eliminated with kinetics comparable to that seen in wild-type crosses (compare Figure 6B with Figure 4).

**Figure 6.**
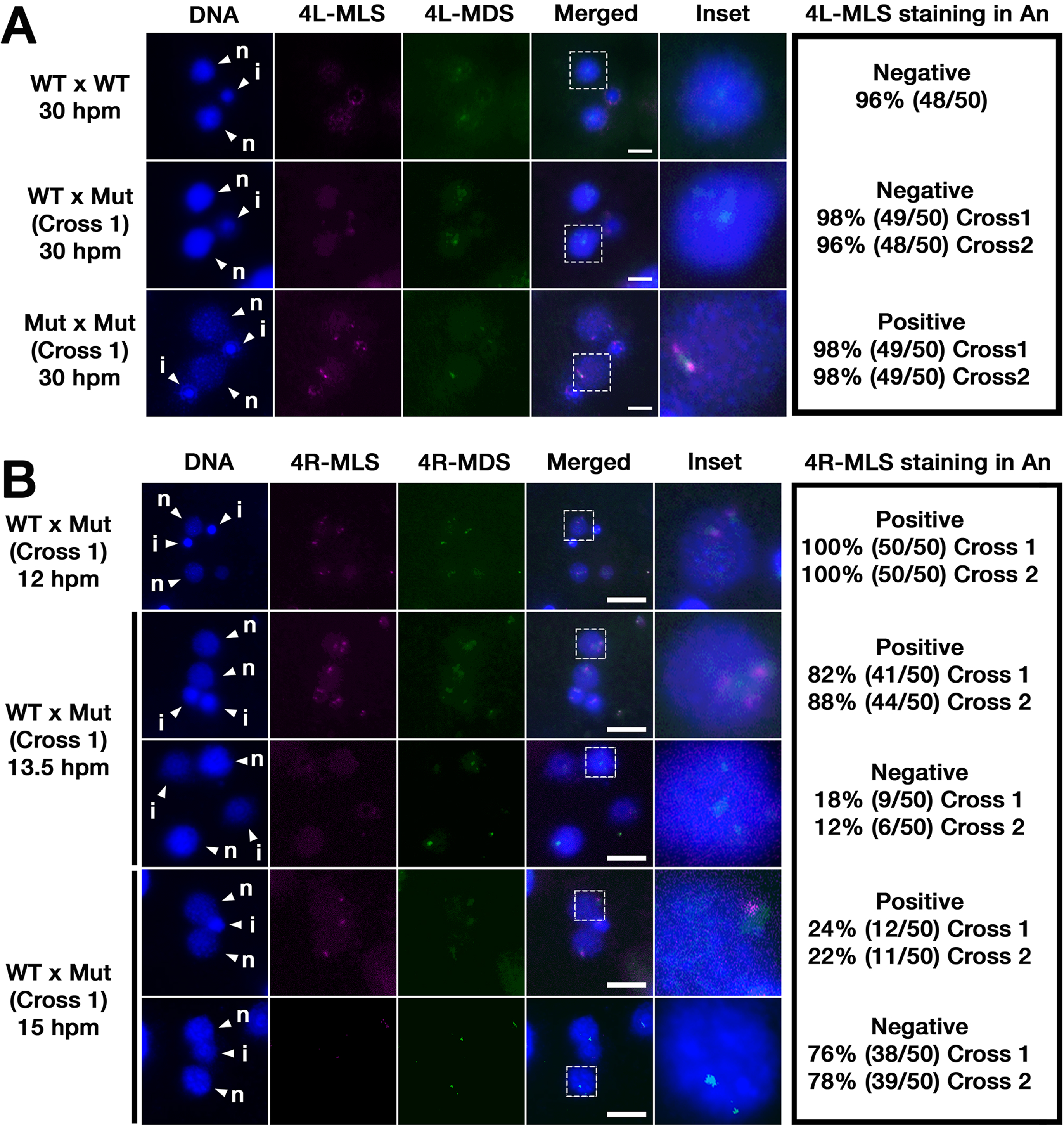
Effect of 4R-CBS mutations on elimination of 4R-MLS. (A) Exconjugants at 30 hours post-mixing (hpm) from crosses between two wild-type strains (WT x WT), two 4R-CBS mutant strains (Mut x Mut), or a wild-type and a 4R-CBS mutant strain (WT x Mut) were analyzed by oligo-FISH using a pool of oligonucleotide probes complementary to 4R-MLS (magenta) and 4R-MDS (green). DNA was counterstained with DAPI (blue). Insets show enlarged views of regions marked by dotted squares. The presence (Positive) or absence (Negative) of the 4R-MLS FISH signal in the developing MAC (An) was examined in 50 cells per cross. Arrowheads with “i” and “n” indicate the MIC and the new MAC, respectively. Scale bars: 10 µm. (B) Exconjugants from a cross between a wild-type strain and a 4R-CBS mutant strain (WT x Mut) at the indicated time points were analyzed as in (A).

We also monitored the behavior of the adjacent 4R-MDS. In wild-type crosses (Figure 7, WT × WT), the number of 4R-MDS foci in the new MAC remained stable from 12 to 13.5 hpm, then increased at 15 hpm as a result of endoreplication. In contrast, in WT × Mut crosses (Figure 7, WT × Mut), the number of 4R-MDS foci dropped transiently at 13.5 hpm before increasing at 15 hpm.

**Figure 7.**
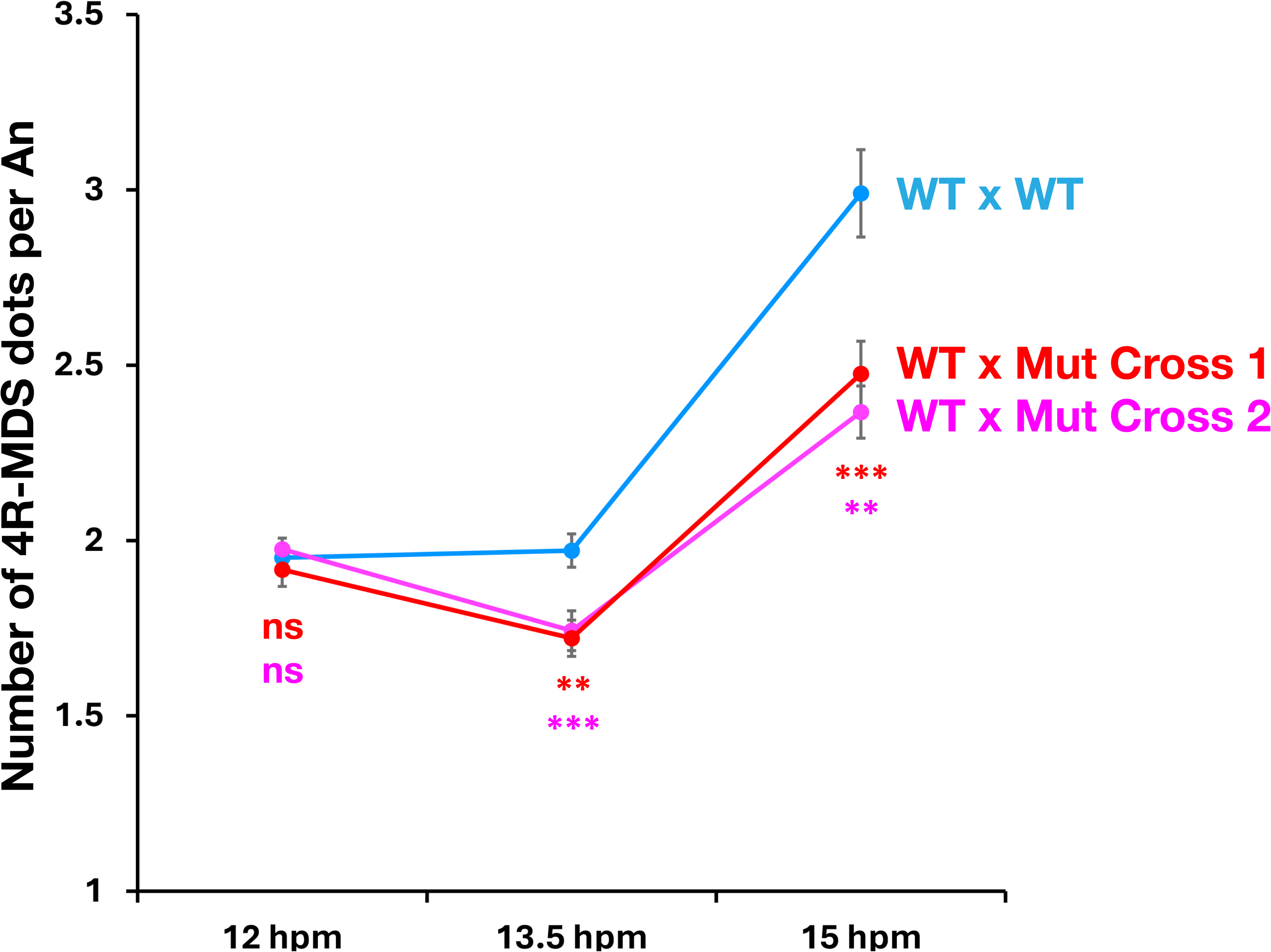
Effect of 4R-CBS mutations on the behavior of 4R-MDS. Exconjugants from a cross between two wild-type strains (WT × WT) and two independent crosses between a wild-type and a 4R-CBS mutant strain (WT × Mut, Cross 1 and Cross 2) were analyzed by oligo-FISH using a pool of oligonucleotide probes complementary to 4R-MDS. The number of 4R-MDS foci per developing MAC (An) at 12 hpm (n = 204), 13.5 hpm (n = 140), and 15 hpm (n = 202) was counted, and the mean ± standard error of the mean (SEM) was plotted. Statistical significance between WT x WT and WT x Mut Cross 1 (red) or Cross 2 (magenta) was assessed by Student’s *t*-test (ns: p > 0.05; **p ≤ 0.01; ***p ≤ 0.001).

These results show that the 4R-CBS mutations do not prevent elimination of 4R-MLS. Instead, they appear to promote elimination of the adjacent 4R-MDS. This likely reflects co-elimination of 4R-MLS and 4R-MDS when chromosome breakage between them is blocked.

### Biparental transmission of 4R-CBS mutations blocks DNA elimination and prevents production of viable sexual progeny

Because 4R-MDS contains 93 predicted genes, its elimination would likely result in the loss of essential functions required for cell viability. Consistent with this idea, no viable sexual progeny were obtained when two 4R-CBS homozygous mutants were crossed (Figure 8A, Mut × Mut). Based on our experimental setup (see Materials and Methods), we estimate that the frequency of viable progeny from these crosses was fewer than 1 per ∼2 x 10⁴ mating pairs. In contrast, viable progeny were produced when the mutants were crossed with wild-type cells (Figure 8A, WT × Mut).

**Figure 8.**
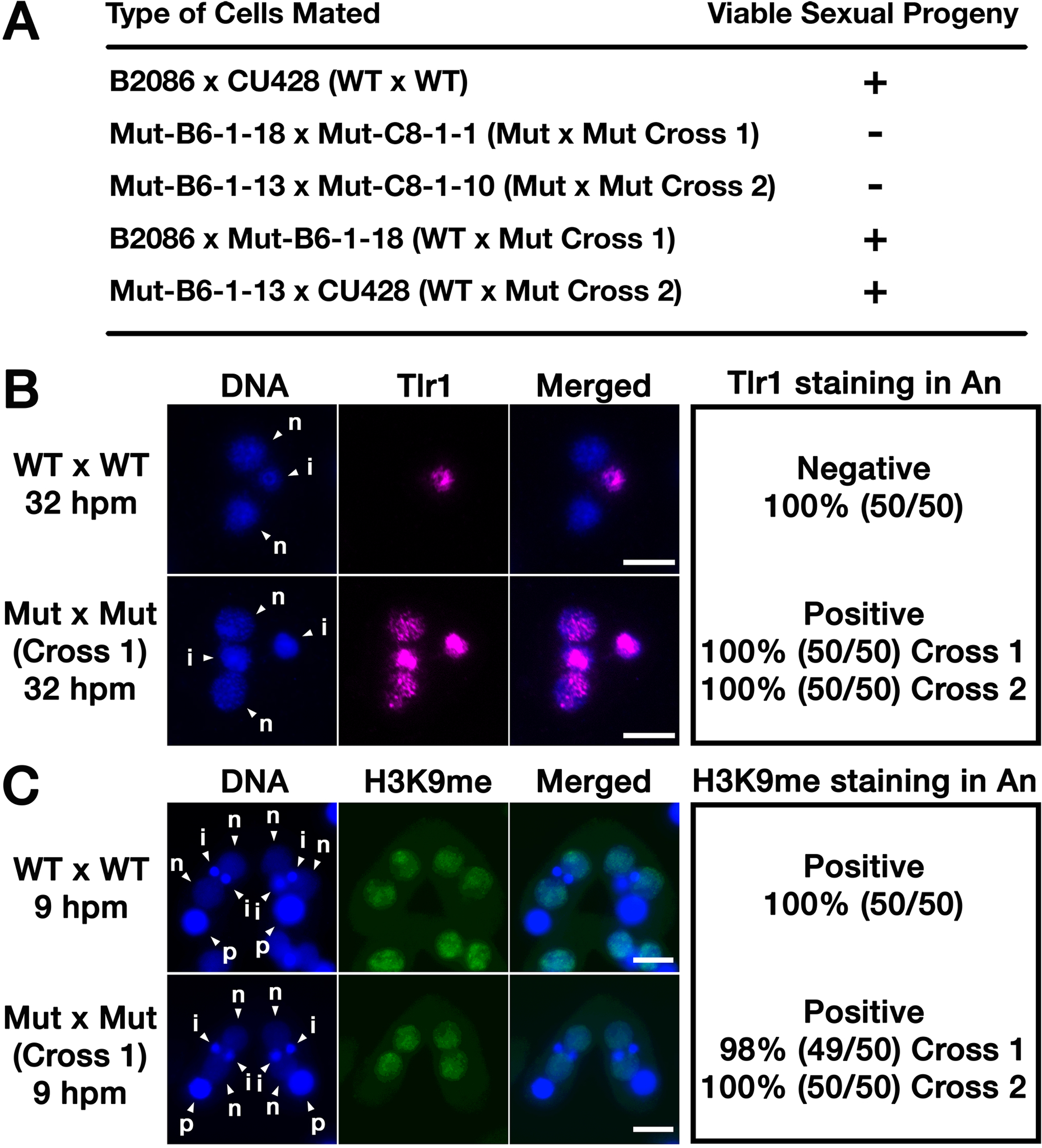
Effect of 4R-CBS mutations on progeny viability, DNA elimination, and heterochromatin formation. (A) The indicated sets of strains were crossed, and the presence (+) or absence (−) of viable sexual progeny was examined. See *Materials and Methods* for details. (B, C) Exconjugants at 32 hours post-mixing (hpm) (B) or conjugating pairs at 9 hpm (C) from a cross between two wild-type strains (WT x WT) and two independent crosses between a wild-type and a 4R-CBS mutant strain (WT x Mut, Cross 1 and Cross 2) were analyzed. FISH was performed using probes complementary to the MIC-specific *Tlr1* element (magenta) (B), and immunofluorescence staining was performed using an antibody against di- and tri-methylated histone H3 at lysine 9 (H3K9me, green) (C). DNA was counterstained with DAPI (blue). The presence or absence of the *Tlr1* FISH signal (B) or H3K9me immunostaining signal (C) in the developing MAC (An) was examined in 50 cells per cross. Arrowheads with “i”, “p”, and “n” indicate the MIC, the parental MAC and the new MAC, respectively. Scale bars: 10 µm.

Unexpectedly, however, instead of promoting co-elimination of 4R-MLS and 4R-MDS, the 4R-CBS mutations led to retention of both regions in the new MAC of exconjugants from Mut × Mut crosses (Figure 6A, Mut × Mut). Further analysis indicated that this was most probably due to a general block of DNA elimination. FISH with Tlr1 probes showed that DNA elimination was disrupted in the new MACs of Mut × Mut exconjugants (Figure 8B). In contrast, immunofluorescence staining with an antibody against di- and trimethylated histone H3 at lysine 9 (H3K9me) indicated that heterochromatin formation in the developing MAC, a prerequisite for DNA elimination (Liu et al., 2007; Taverna et al., 2002; Xu et al., 2021), was not obviously impaired (Figure 8C).

Taken together, these results indicate that physical separation of MLSs and MDSs by chromosome breakage, at least at the right arm of chromosome 4, is required for proper programmed DNA elimination and the formation of viable sexual progeny.

### Biparental transmission of 4R-CBS mutations does not specifically disrupt gene expression in the proximal MDS

We hypothesized that chromosome breakage at the 4R-CBS prevents the spread of heterochromatin into the adjacent 4R-MDS, thereby preserving expression of genes within this region, including those required for DNA elimination.

To test this hypothesis, we compared mRNA expression profiles in Mut × Mut crosses with those in WT crosses by RNA-seq at 13.5 hpm and 15 hpm, when chromosome breakage and 4R-MLS elimination are, respectively, ongoing and nearly complete in WT cells. Contrary to our expectation, mRNA expression of most genes encoded within the 4R-MDS (MAC chromosome 164) was unaffected in Mut × Mut crosses (Figure 9A). Moreover, these genes were not significantly more affected than genes across the rest of the genome (Figure 9B).

**Figure 9.**
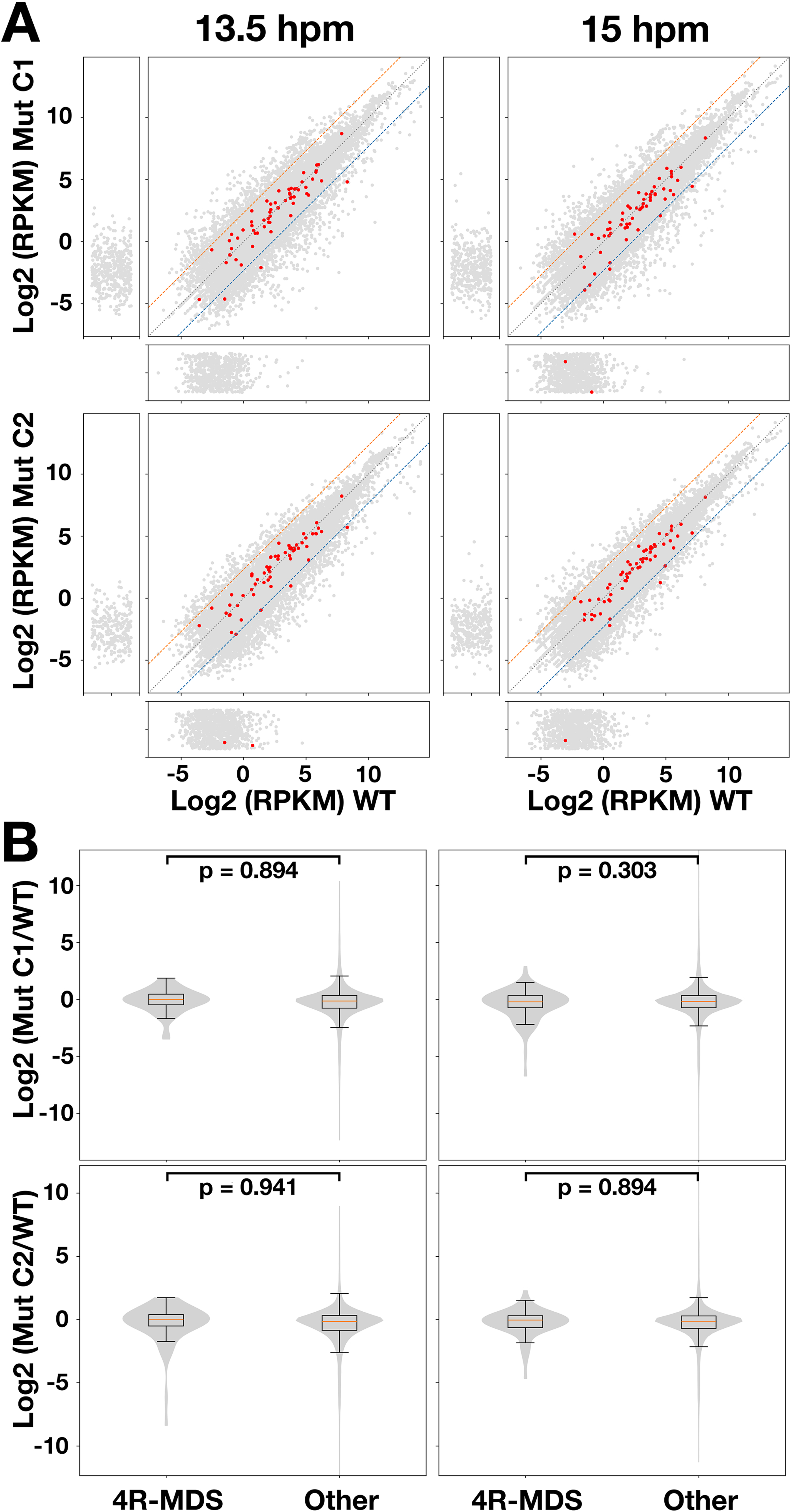
Effect of 4R-CBS mutations on gene expression from 4R-MDS. (A) Scatter plots show log₂-transformed gene expression levels for individual genes in wild-type (WT, x-axis) versus two independent crosses of 4R-CBS mutants (Mut C1 and C2, y-axis) samples at 13.5 hpm (left) and 15 hpm (right). Expression values were thresholded at a minimum of 0.005 prior to log₂ transformation and plotted up to 20,000 normalized units. Each point represents one gene; genes encoded in 4R-MDS are highlighted in red. The marginal side panels show genes with expression below the threshold (<0.005) in either WT or Mut. Dashed orange and blue lines respectively represent +5- and -5-fold differential expression boundaries. (B) Box and violin plots show the distribution of gene expression changes between 4R-CBS mutant (Mut C1 or C2) and wild-type cells (WT) at 13.5 hpm (left) and 15 hpm (right) for genes encoded in 4R-MDS compared with all other genes. Boxes indicate the interquartile range, center lines indicate medians, and whiskers extend to 1.5x the interquartile range. A Mann–Whitney U test was used to compare the distributions of the two gene groups, and the resulting p-values are shown.

These results indicate that the global DNA elimination defect caused by biparental transmission of 4R-CBS mutations is unlikely to result from erroneous heterochromatin spreading into the adjacent MDS.

## Discussions

In this study, we investigated MIC telomere elimination during MAC development in *Tetrahymena*. Using optimized oligo-FISH, we tracked the MIC-specific G_4_T_3_ repeat and found telomere elimination occurs alongside IES removal (Figure 2). Genome comparisons indicated that MIC telomeres and adjacent MLSs are co-eliminated via chromosome breakage (Figure 3). Visualization of MLSs confirmed elimination with timing similar to MIC telomeres (Figure 4). CRISPR–Cas9 disruption of the 4R-CBS showed that 4R-MLS elimination was not blocked but caused loss of the adjacent 4R-MDS (Figures 6, 7), suggesting their co-elimination when breakage is impaired. In biparental mutant crosses (Mut × Mut), both 4R-MLS and 4R-MDS were retained, DNA elimination was broadly disrupted, and no viable progeny were produced (Figure 8). Thus, chromosome breakage at CBSs is essential for separating MLSs from MDSs, ensuring proper DNA elimination and sexual progeny development.

Our data support a critical role for 4R-CBS in separating 4R-MLS from 4R-MDS, but it remains unclear whether all MIC chromosome ends are strictly CBS-dependent for their elimination. We found no conserved CBS sequence at two of ten MDS–MLS borders (Figure 3A). This may reflect CBS-independent processing or assembly errors in repetitive MLS regions that obscure CBSs. Clarifying this will require re-examining MLS–MDS borders and systematically mutating MLS-proximal CBSs across chromosome ends. So far, attempts to mutate other CBSs have failed, as the AT-rich *Tetrahymena* genome limits Cas9 targeting of the 15-nt CBS. Thus, alternative genome-editing tools will be essential to dissect CBS function at MIC chromosome ends.

The parallel elimination of MIC telomeres, MLSs, and IESs suggests the existence of a mechanism that synchronizes these processes. While blocking chromosome breakage could trigger checkpoint-like responses that prevent progression of DNA elimination, this is inconsistent with the observation that uniparental inheritance of the 4R-CBS mutation did not inhibit sexual progeny formation (Figure 8A), which requires DNA elimination (Cheng et al., 2010). Moreover, chromosome breakage can be inhibited without disrupting DNA elimination, as shown in cells lacking zygotic expression of the p68-like RNA helicase Drh1 (McDaniel et al., 2016).

Instead, coordination likely stems from a shared mechanism. We previously showed that MIC chromosomal ends, like IESs, produce abundant ∼29-nt scnRNAs (Hamilton et al. 2016; Noto and Mochizuki 2017) that act with Twi1 to promote Polycomb-dependent heterochromatin formation and DNA elimination (Mochizuki et al., 2002; Noto et al., 2015; Xu et al., 2021). In addition, telomere-complementary scnRNAs were reported to be produced specifically during conjugation (Cao et al., 2009). Thus, MIC telomere and MLS elimination are likely driven by the same scnRNA-directed pathway as IESs, providing a basis for their coordination.

The 4R-CBS mutation caused different phenotypes depending on its uniparental or biparental inheritance (Figures 6–8). When inherited uniparentally (WT × Mut cross), 4R-MLS and 4R-MDS were co-eliminated, likely because blocking the chromosome break that normally separates these regions causes them to co-segregate into the nuclear compartment where DNA elimination occurs. In contrast, when inherited biparentally (Mut × Mut cross), DNA elimination was globally impaired.

Our gene expression analysis revealed that genes within 4R-MDS were not specifically affected in Mut x Mut crosses relative to WT, making it unlikely that chromosome breakage at the 4R-CBS normally functions as an insulator that blocks heterochromatin spreading into the 4R-MDS and preserves gene expression in this region. Instead, inhibition of chromosome breakage at 4R-CBS, or the mutation at this site itself, may cause architectural defects that compromise global DNA elimination.

Previous studies have shown that MIC chromosomes exhibit TAD-like structures, with boundaries occurring at CBSs, and have proposed that these domains may contribute to MAC chromosome formation (Luo et al., 2020). The 4R-CBS mutation could disrupt these TAD-like structures in the MIC, potentially affecting either scnRNA production during early conjugation or the chromatin architecture required for DNA elimination in the developing MAC at later stages of conjugation.

The presence of MLSs at all MIC chromosome ends indicates that there must be a biological requirement for MLSs at each end. Similarly, some nematodes also undergo programmed DNA elimination in somatic cells during embryogenesis and eliminate telomeres together with germline-limited sequences through chromosome breakage at all germline chromosome ends (Dockendorff et al., 2022; Estrem et al., 2024; Simmons et al., 2024). Therefore, co-elimination of telomeres and large blocks of germline-limited sequences is a common strategy, probably by convergent evolution, for ensuring switching germline to somatic telomeres through programmed DNA elimination in ciliates and nematodes. The observed link between chromosome breakage at 4R-CBS and the essential DNA elimination process may reflect the biological significance of MLSs and the importance of their removal from the MAC. Coupling these processes may have evolved as a mechanism to ensure that only functional chromosome-end CBS loci are preferentially transmitted to future generations.

Because *Tetrahymena* strains lacking some MIC chromosomal arms including MLSs can be established (Bruns et al., 1983, 1982), MLSs are likely to be unnecessary for maintenance of MIC chromosomes during vegetative growth but act during conjugation. Besides ensuring MIC telomere elimination, MLSs may serve as scnRNA sources that target transposon-derived sequences, similar to piRNA clusters in animals where transposon remnants generate piRNAs (Czech et al., 2018; Huang et al., 2017; Iwasaki et al., 2015). Their adjacency to telomeres may be required because telomeres could act as promoters of non-coding RNA transcriptions producing scnRNAs during early conjugation, consistent with G/C-rich tracts being enriched at transcription start sites of scnRNA precursors (Cai et al., 2026, 2025). Future studies should address the biological significance of MLSs by specifically deleting MLSs from the MIC. Such targeted deletions may also result in retention of the adjacent MIC telomere in the MAC, allowing investigation of whether the presence of MIC telomeres in the MAC affects cell physiology.

## Materials and methods

### Strains and Culture Conditions

Cells were cultured at 30 °C in 1× SPP medium (Gorovsky et al., 1975) containing 2% (w/v) proteose peptone. For induction of conjugation, log-phase cultures (∼5-7 × 10⁵ cells/mL) of complementary mating types were washed once in 10 mM Tris-HCl (pH 7.5), starved in the same buffer for 12-24 h, and then mixed at a 1:1 cell ratio to a final concentration of 5 × 10⁵ cells/mL at 30 °C.

### Probe design and production for Fluorescent in situ hybridization (FISH)

To detect the MIC-specific G_4_T_3_ telomere repeat, an oligonucleotide consisting of five tandem repeats of 5’-CCCCAAA-3’, flanked by FLAP-X complementary sequences at both ends, was synthesized by Eurofins Genomics. Cy3-labeled probes for the Tlr1 element were produced by nick translation as described previously (Noto et al., 2010). To design oligo probes that uniquely recognize the 3L-MLS, 4R-MLS, or 4R-MDS genomic regions, all possible continuous 32-nt sequences were extracted from each region. Sequences unaligned to the MIC genome (PRJCA042635, Genome Warehouse database, China National Center for Bioinformation), which lacks the 3L-MLS, 4R-MLS, or 4R-MDS regions, were selected using Bowtie2 (sensitive-local option). Sequences with low (<40%) or high (>70%) GC content were removed, and 384 non-overlapping oligonucleotides were manually selected. For probes targeting 3L-MLS and 4R-MLS, the sequence complementary to FLAP-X was added to the 5’ ends. For probes targeting 4R-MDS, the sequence complementary to FLAP-Y was added to the 5’ ends. Each pool of 384 oligos was synthesized by Integrated DNA Technologies (oPools Oligo Pools) and dissolved in TE buffer at 100 µM. FLAP-X and FLAP-Y oligos used as secondary probes were described previously (Tsanov et al., 2016). FLAP-X and FLAP-Y oligonucleotides, modified with Cy3 and Cy5 at both ends, respectively, were synthesized by Integrated DNA Technologies. All oligonucleotide sequences used in this study are listed in Supplementary Table S2 and S3.

### FISH

Approximately 3 × 10⁷ cells were pelleted by centrifugation at 400 × g for 3 min, resuspended in Carnoy’s fixative (methanol/acetic acid, 3:1), and incubated for 1 h at room temperature (RT). Cells were then pelleted again at 400 × g for 2 min, resuspended in 0.6 mL Carnoy’s fixative, and 50 µL was spread onto Superfrost Plus Adhesion Microscope Slides (Epredia, J7800AMNZ). Slides were air-dried and stored at –20 °C until use.

Slides were equilibrated to RT for 3 min and rehydrated in water at RT for 5 min. They were then incubated in RIPA+ buffer (150 mM NaCl, 2% NP-40, 1% sodium deoxycholate, 0.2% SDS, 1 mM EDTA [pH 8.0], 50 mM Tris-HCl [pH 8.0]) at 30 °C for 1 h. The buffer was replaced with fresh RIPA+ buffer, and incubation continued for an additional 1 h at 30 °C. Next, slides were washed twice in 30 mM Tris-HCl (pH 8.0) for 5 min each, then incubated in Proteinase K solution (0.2 µg/mL Proteinase K [Macherey-Nagel] in 30 mM Tris-HCl [pH 8.0]) at 30 °C for 30 min. This was followed by two washes in 0.2% glycine in PBS for 5 min each, and two washes in PBS for 5 min each. Slides were then incubated twice with RNase A solution (0.1 mg/mL RNase A [Thermo Fisher Scientific] in PBS containing 0.1% Tween-20 [PBST]) at 30 °C for 30 min each, followed by washes in PBST for 10 min and PBS for 5 min. Finally, slides were post-fixed in 70% ethanol and air-dried.

After rehydration in water for 2 min, slides were incubated in 70% formamide, 2× SSC at 70 °C for 2 min, and immediately rinsed in ice-cold water for 2 min. Hybridization buffer (50% formamide, 10% dextran sulfate, 2× SSC) containing 1.25 µM primary probes (for 3L-MLS, 4R-MLS, and 4R-MDS), 0.25 µM probes (for G_4_T_3_), or 0.5 ng/µL probes (for Tlr1) was applied to slides. Coverslips were sealed with rubber cement, slides were incubated at 80 °C for 10 min, and then incubated overnight at 37 °C. Slides were washed twice in 50% formamide, 2× SSC, 0.1% SDS at 37 °C for 10 min each, once in 2× SSC, 0.1% SDS at 37 °C for 10 min, and once in 0.5× SSC at RT for 5 min. Next, slides were incubated in hybridization buffer containing 1 µM FLAP-X–Cy3 and/or FLAP-Y–Cy5 for 2 h at 30 °C. Slides were then washed twice in 50% formamide, 2x SSC at 37 °C for 10 min each, once in 2x SSC at 37 °C for 10 min, and once in 0.5× SSC at RT for 5 min. For Tlr1-FISH, the incubation and washing steps with secondary probes were omitted. Cells were counterstained with 10 ng/mL 4’, 6-diamidino-2-phenylindole (DAPI; Thermo Fisher Scientific) in 0.05% Triton X-100, PBS for 10 min, followed by washing in 0.05% Triton X-100, PBS for 10 min. Samples were mounted with ProLong Gold Antifade Reagent (Thermo Fisher Scientific).

Images were acquired using a Zeiss Axio Imager Z2 equipped with a Plan-Apochromat 100×/1.4 Oil M27 objective, filter sets 49 (DAPI), 43 HE (Cy3), and 50 (Cy5), and 353, 545, and 650 nm excitation lasers with 150 ms exposure. Fifty cells were examined at each cross and each time point.

### Identification and analysis of MLS regions at the ends of MIC chromosomes

The *Tetrahymena thermophila* MIC genome sequence, in which five MIC chromosomes are assembled at a telomere-to-telomere level (PRJCA042635, Genome Warehouse database, China National Center for Bioinformation), was compared with the *Tetrahymena thermophila* MAC genome sequence (assembly_v5, available at https://tet.ciliate.org/). The MAC genome sequence was aligned against the MIC genome using Minimap2 (Galaxy Version 2.28) (Li, 2018) with default parameters. Mapping results were converted into a coverage bigWig file using the bamCoverage tool (Galaxy Version 3.5.4) within deepTools2 (Ramírez et al., 2016) and visualized with the Integrative Genomics Viewer (Version 2.13.0).

To assess the repetitiveness of each MLS, all possible 20-nt sequences from the MIC genome were mapped back to the same MIC genome using HIAST2 (Galaxy Version 2.2.2). The number of uniquely mapped reads within each MLS region was then determined using Samtools view (Galaxy Version 1.22) with a quality threshold of 5. Regions lacking coverage by uniquely mapped reads were classified as repetitive.

### Construction of 4R-CBS mutants

The most proximal CBS at the right arm of the MIC chromosome 4 (4R-CBS) was mutated using a CRISPR-Cas9 system adapted for *Tetrahymena* (Suhren et al., 2017). To construct the 4R-CBS editing plasmid, 4R-Cas9T1-S and 4R-Cas9T1-AS (Supplementary Table S3) were annealed and ligated into the BbsI site of the pC9T vector. The resulting plasmid was digested with XhoI and introduced into the MAC *BTU1* locus of the B2086 and CU428 strains by homologous recombination using biolistic transformation (Bruns and Cassidy-Hanley, 2000). Transformants were selected in 100 µg/mL paromomycin (Sigma–Aldrich). Three transformants per strain were selected and subjected to phenotypic assortment as described previously (Hamilton and Orias, 2000) until the cells were able to grow in 1 mg/mL paromomycin. Then, Cas9 expression was induced by incubating cells in 1 µg/mL CdCl2 in 1 x SPP for 6 h, and single cells were isolated into SPP drops. Twenty-four clones from each transformant line were recovered in SPP medium in 96-well plates, and the 4R-CBS locus was examined by direct cell PCR using primers 4R-2021T2T-Mut-cFW and 4R-2021T2T-Mut-cRV (Supplementary Table S3), followed by sequencing the PCR products. Three and two heterozygous mutants derived from B2086 and CU428, respectively, were obtained. Finally, to obtain homozygous 4R-CBS mutants, Round I genomic exclusion was induced by crossing the heterozygous mutants with strain B*VI. The resulting Round I exconjugants were either homozygous mutant or wild-type for the 4R-CBS locus. Homozygous 4R-CBS mutants were identified by direct cell PCR followed by Sanger sequencing. Obtained two homozygous 4R-CBS mutants from B2086 (B6-1-13 and B6-1-18) and two from CU428 (C8-1-10 and C8-1-1) were used for further studies.

### Detection of *de novo* telomere formation

Genomic DNA was extracted as previously described from overnight-starved cells, mating cells, and/or exconjugants at different time points. A total of 10 ng of genomic DNA was used for PCR with the primers Tel2 and 4R-Tel-1 for 4R-MLS, or Tel2 and 3L-Tel-2 for 3L-MLS (25 µL reaction volume, using Taq DNA polymerase [NEB, M0273]; 28 cycles of 95 °C for 20 s, 53 °C for 30 s, and 68 °C for 1 min). Then, 0.5 µL of the PCR product was used as a template for a second PCR with the primers Tel and 4R-Tel-2 for 4R-MLS, or Tel and 3L-Tel-3 for 3L-MLS (under the same conditions as the first PCR).

### Viability test of sexual progeny

Two strains with complementary mating types were individually starved overnight in 10 mM Tris at a concentration of ∼5 × 10^5^ cells/mL and then mixed to induce conjugation. At 24-27 hpm, 100 µL of culture was combined with 900 µL of 1 x SPP in a 24-well plate and incubated for 3 h. Then, 6-methylpurine (6-mp, Sigma–Aldrich) was added to a final concentration of 15 µg/mL and incubated for 4 days. Because all crosses used in this study used CU428 or CU428-derived strains, which are homozygous for a 6-mp resistance mutation in their MIC, successful completion of conjugation should result in the production of 6-mp-resistant sexual progeny. The presence of 6-mp-resistant progeny was examined under a dissection microscope. Each cross was repeated three times to confirm reproducibility.

### 4R-MLS and 4R-MDS detection in 4R-CBS mutants

The presence of 4R-MLS and 4R-MDS was examined by oligo-FISH as described above. Cells were fixed at 30 hpm, and 50 exconjugants per cross were analyzed. For crosses between wild-type cells and 4R-CBS mutants (WT × Mut cross), cells were additionally fixed at 12, 13.5, and 15 hpm and analyzed. An exconjugant was considered 4R-CBS positive if one or both new MACs exhibited 4R-MLS FISH staining. The number of 4R-MDS foci in 204, 140, and 202 new MACs from the WT × Mut cross at 12, 13.5, and 15 hpm, respectively, was counted and statistically analyzed using Student’s t-test.

### Immunofluorescence staining

Approximately 3 × 10⁷ mating cells were fixed at 9 hpm and permeabilized in 3.7% formaldehyde, 0.5% Triton X-100 at RT for 30 min. After centrifugation, the cells were resuspended in 1 mL 3.7% formaldehyde containing 3.4% sucrose, and 50 µL of the suspension was spread on a slide and air-dried. Slides were stored at –20°C until use. Slides were rehydrated in water for 5 min and then incubated with blocking buffer (0.1% Tween 20, 3% bovine serum albumin, 10% normal goat serum in PBS) at RT for 2 h. After washing in PBST, slides were incubated overnight at 4°C with monoclonal anti-H3K9me2/3 antibody (6F12, Cell Signaling) diluted 1:250 in blocking buffer. Following three 10-min washes in PBST, slides were incubated at RT for 1 h with Alexa Fluor 488-conjugated goat anti-mouse IgG (Invitrogen, A-1100) diluted 1:1000. Slides were washed in PBST for 10 min, incubated with 10 ng/mL DAPI in PBST for 10 min, and washed again for 10 min. Cells were mounted with Prolong Gold Antifade Reagent. Fifty cells were analyzed per cross at each time point.

### RNA-seq

Total RNA was extracted from cell pellets using TRIzol reagent, followed by purification with the RNA Clean & Concentrator™-5 kit (Zymo Research) according to the manufacturer’s protocol, including on-column DNase I treatment to remove genomic DNA contamination. RNA was eluted in nuclease-free water, and RNA concentration was measured using the Qubit RNA Broad Range (BR) Assay Kit (Invitrogen). Samples were submitted to Azenta Life Sciences for poly(A)+ RNA selection, library preparation, and paired-end sequencing on an Illumina NovaSeq 6000 platform (PE150). Raw sequencing reads are available in the European Nucleotide Archive (ENA) under accession numbers ERR17144311–ERR17144316.

Low-quality bases and adapter sequences were trimmed. The processed reads were aligned to the *Tetrahymena thermophila* MIC genome described above using HISAT2 (Galaxy version 2.2.2) with default parameters for paired-end mapping. Reads that mapped to the MIC genome were then aligned to the *Tetrahymena thermophila* gene model (v6, available at https://tet.ciliate.org/) using HISAT2. The resulting alignment files in SAM format were converted to binary (BAM) format, coordinate-sorted using Samtools view (Galaxy version 1.22), and indexed using Samtools idxstats (Galaxy version 2.0.8).

## Acknowledgements

We are deeply grateful to Jie Xiong, Wei Miao (Chinese Academy of Sciences) and Yifan Liu (University of Southern California) for sharing unpublished MIC genome assembly data, which was essential for the analyses presented in this paper. We acknowledge the MRI facility, member of the national infrastructure France-BioImaging supported by the French National Research Agency (ANR-10-INBS-04). This work was supported by Equipes a FRM 2022 grant from the Foundation Recherche Médicale (FRM, EQU202203014651), an ARC 2021 PJA3 grant from the ARC Foundation (ARCPJA2021060003830), and a PRC Grant, from the French National Research Agency (ANR, ANR-24-CE12-3978-01) to KM.

## Figure Legends

**Supplementary Figure S1. Elimination of Tlr1 element during MAC development**

Conjugating wild-type cells at the indicated time points (hours post-mixing, hpm) were analyzed by FISH using probes complementary to the MIC-specific Tlr1 element (magenta). DNA was counterstained with DAPI (blue). The Tlr1 FISH signal in the developing MAC (An) was examined in 50 cells per time point and classified according to staining pattern: Homogeneous, signal distributed throughout the new MACs; DE-body, signal localized to peripheral foci (DNA elimination bodies); or No staining, no detectable signal. Scale bars: 10 µm.

**Supplementary Figure S2. Elimination of 3L-MLS during MAC development in wild-type cells**

Conjugating wild-type cells at the indicated time points (hours post-mixing, hpm) were analyzed by oligo-FISH using pool of oligonucleotide probes complementary to 3L-MLS (magenta). DNA was counterstained with DAPI (blue). Insets show enlarged images of the regions indicated by dotted squares. The presence (Positive) or absence (Negative) of the 3L-MLS FISH signal in new MAC (An) in 50 cells per time point was examined. Scale bars: 10 µm.

