## Supplementary Figures and Tables for "Linking Germline Telomere Removal to Global Programmed DNA Elimination in *Tetrahymena* Genome Differentiation": Supplementary Figures.pdf

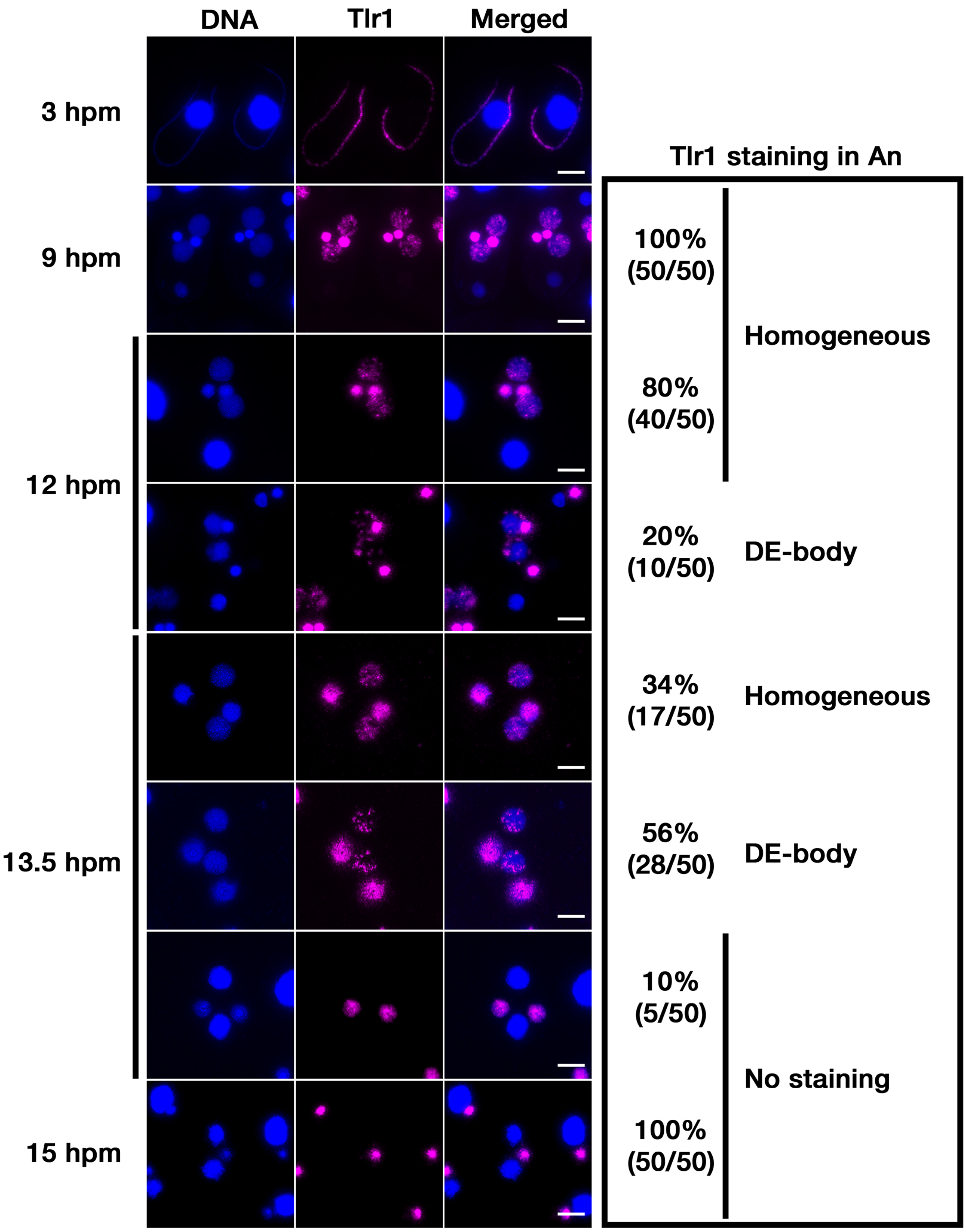


**Supplementary Figure S1. Elimination of Tlr1 element during MAC development**

Conjugating wild-type cells at the indicated time points (hours post-mixing, hpm) were analyzed by FISH using probes complementary to the MIC-specific Tlr1 element (magenta). DNA was counterstained with DAPI (blue). The Tlr1 FISH signal in the developing MAC (An) was examined in 50 cells per time point and classified according to staining pattern: Homogeneous, signal distributed throughout the new MACs; DE-body, signal localized to peripheral foci (DNA elimination bodies); or No staining, no detectable signal. Scale bars: 10 µm.

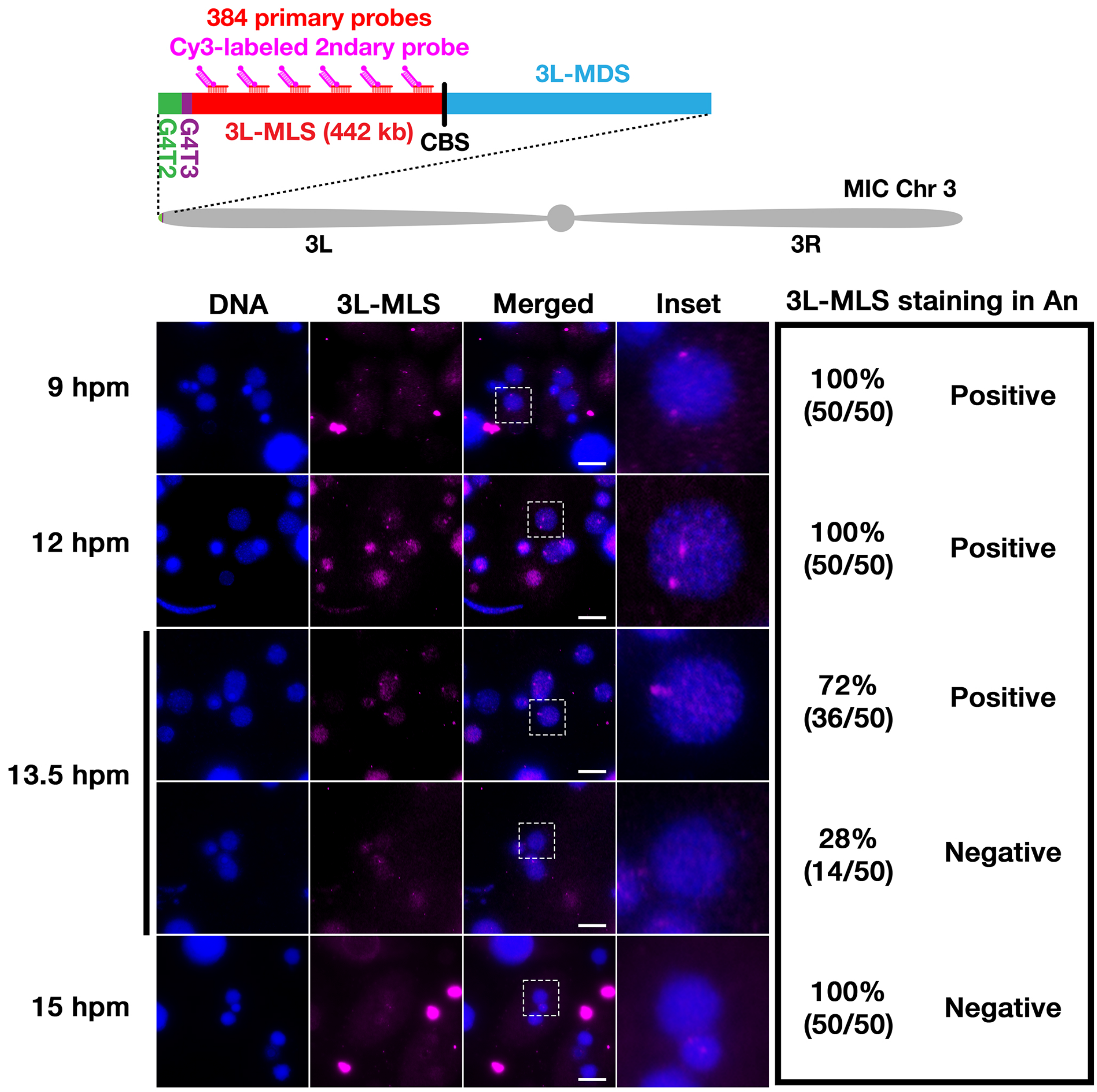


**Supplementary Figure S2. Elimination of 3L-MLS during MAC development in wild-type cells**

Conjugating wild-type cells at the indicated time points (hours post-mixing, hpm) were analyzed by oligo-FISH using pool of oligonucleotide probes complementary to 3L-MLS (magenta). DNA was counterstained with DAPI (blue). Insets show enlarged images of the regions indicated by dotted squares. The presence (Positive) or absence (Negative) of the 3L-MLS FISH signal in new MAC (An) in 50 cells per time point was examined. Scale bars: 10 µm.
