## Supplementary figures and images for "Linking Germline Telomere Removal to Global Programmed DNA Elimination in *Tetrahymena* Genome Differentiation"

### Supplementary Table_S1.docx

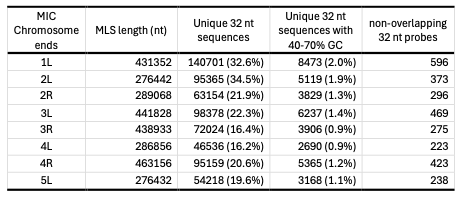

**Supplementary Table S1. Designing MLS-specific Oligo Probes**
